# 300 years of change for native fish species in the upper Danube River Basin – historical flow alterations versus future climate change

**DOI:** 10.1101/2021.06.14.448400

**Authors:** Martin Friedrichs-Manthey, Simone D. Langhans, Florian Borgwardt, Thomas Hein, Harald Kling, Philipp Stanzel, Sonja C. Jähnig, Sami Domisch

**Author notes:** Authors contributed equally (Sonja C. Jähnig, Sami Domisch). corresponding author: Martin Friedrichs-Manthey, Mail.

## Abstract

River ecosystems belong to the most threatened ecosystems on Earth. Historical anthropogenic alterations have, and future climate change will further affect river ecosystems and the species therein. While many studies assess the potential effects of expected future changes on species, little is known about the severity of these changes compared to historical alterations. Here, we used a unique 300-year time series of hydrological and climate data to assess the vulnerability of 48 native fish species in the upper Danube River Basin. We calculated species-specific vulnerability estimates relative to the reference period (1970-2000) for the periods 1800-1830, 1900-1930, and 2070-2100, including two Representative Concentration Pathways (RCP 4.5 and 8.5) and identified the environmental drivers of vulnerability estimates. Models showed that future changes under RCP 4.5 would result in moderate species vulnerability compared to historical conditions, while under RCP 8.5, the vulnerability for all species increased substantially. In addition, species vulnerability was mainly driven by hydrology in the past and is likely to be driven by temperature in the future. Our results show that future climate change would alter environmental conditions for riverine fish species at a similar magnitude as historical anthropogenic hydrological river alterations have. Shedding light on such long-term historical and possible future anthropogenic alterations provides valuable insights for prioritising conservation actions for riverine fish species.

## Introduction

Freshwaters belong to the most threatened ecosystems on Earth (Dudgeon, 2019; Reid et al., 2018), with almost one-third of all freshwater species facing the threat of extinction (Collen et al., 2014) and a global freshwater vertebrate population decline of 84% on average in the past 50 years (WWF, 2016). Among all known freshwater fish species, one quarter faces extinction (Su et al., 2021). Fish species in river ecosystems react sensitively to alterations in discharge (Beatty et al., 2014; Rolls & Arthington, 2014; Ward et al., 2015; Xenopoulos & Lodge, 2006) and in temperature (Buisson et al., 2008; Buisson & Grenouillet, 2009; Comte et al., 2013; Kriaučiūnienė et al., 2019; Lyons et al., 2010). Globally riverine fish species suffered in the past mainly from anthropogenic discharge alterations (Böhm & Wetzel, 2010; Grill et al., 2019; Haidvogl et al., 2014; Hohensinner et al., 2004), while pronounced long-term temperature increases were only relatively recently observed (IPCC, 2017). In contrast, for the future, studies suggest that the significant rise in air temperatures and changes in precipitation patterns (with consequential but probably less pronounced changes in discharge) will be the main driver of vulnerability, i.e. susceptibility to being negatively affected by climate change (Pacifici et al., 2015), for riverine species (Jaric et al., 2019; Kriaučiūnienė et al., 2019; Reid et al., 2018).

The majority of large rivers globally have been modified by humans over decades to meet social demands such as transportation, energy production, flood and disease control, or drinking and agricultural water supply (Grill et al., 2019; Grizzetti et al., 2017; Jungwirth et al., 2014), resulting in a severe loss of natural characteristics of rivers (Cazzolla Gatti, 2016; Wohl, 2019). Climate change scenarios predict a further, significant increase in pressures for river ecosystems within the near future (Dudgeon, 2019; Grill et al., 2019; Jaric et al., 2019; Rodell et al., 2018). For example, climate change will result in increased water temperatures (IPCC, 2017), which often results in a reduction of suitable habitats for native species (Markovic et al., 2014) or an increase in thermal stress as species will be subjected to their upper thermal boundaries (Crear et al., 2020; Till et al., 2019). In addition, expected increases in water use and changes in the amount and the spatial distribution of precipitation (Rodell et al., 2018) will result in enhanced hydrologic pressures for biotic communities in rivers (Kakouei et al., 2018; Rolls et al., 2017; Yoshikawa et al., 2014). One of the many rivers with a long history of human alteration and expected severe climate change effects in the future is the Danube River, one of the largest and most fish species divers rivers in Europe (Jungwirth et al., 2014). For the Danube River Basin the 19^th^ century was dedicated to flood prevention and channelization, especially for the upper part (Jungwirth et al., 2014). However, severe impacts on the fish fauna started at the end of the 19^th^ century, when channelization reached its maximum and soon after the first hydroelectric power stations were built (Jungwirth et al., 2014; Zauner & Schiemer, 1994). In 1956 the first hydroelectric power station in the main channel was completed, and to date, more than 70 hydroelectric power stations exist in the main stem of the upper Danube River (Jungwirth et al., 2014). Considering future alterations, mean annual temperature is predicted to steadily increase with an accelerating rate (IPCC, 2017) in the upper Danube River Basin (Jacob et al., 2014). However, the predictions for changes in precipitation are more divers and effects on the fish fauna are difficult to anticipate (Giorgi et al., 2016). A reduction in precipitation, especially in the summer months, is expected, but some regional climate models also predict an increase with a change from rain to snow, especially in higher alpine areas (Giorgi et al., 2016).

When assessing conservation needs for freshwater biodiversity under future climate change scenarios, the majority of studies neglect the often dramatic historical environmental alterations (Wohl, 2019) and their impact on species or populations. Without quantifying historical alterations, with “historical” referring to a time period in which major anthropogenic changes started until the current point in time, and their impact on species, future predictions can only deliver relative estimates of vulnerability, i.e., relative to the current point in time. Studies focussing on future predictions of possible impacts on species or populations (Kakouei et al., 2018; Markovic et al., 2017; McMahan et al., 2020; Radinger et al., 2017), deliver certainly crucial information for effective conservation planning and management (Bonebrake et al., 2018; Domisch et al., 2019), however predictions often come with high uncertainty (Yates et al., 2018) and are likely to e.g. overestimate the importance of temperature, because the expected rapid increase in temperature will be more pronounced than a gradual change in discharge driven by precipitation changes. Such uncertainties can hinder practical implication of modelling results (McShea, 2014; Porfirio et al., 2014; Schuwirth et al., 2019). Therefore, placing predicted future changes into a historical context can deliver useful information regarding the magnitude of change that species have already been exposed to, compared to what they would expect under future climate change scenarios. Such information would significantly increase the effectiveness of current conservation management efforts (Bonebrake et al., 2010; Novaglio et al., 2020; Pont et al., 2015).

Here, we employ a unique historical time series of observed (1800-2007) and modelled (2007-2100) climate and hydrological data for a 300-year period from 1800 to 2100 for the area of the upper Danube River Basin (Fig. 1). The extensive period allows comparing the effects of major historical alterations in discharge and temperature on the vulnerability of fish species (Jungwirth et al., 2014; Zauner & Schiemer, 1994) with predicted effects for the near future driven by expected alterations in climate conditions (Klein et al., 2011; Kling, Fuchs, et al., 2012; Stanzel & Kling, 2018). In addition, the time series allows quantifying the drivers of expected vulnerability for the historical alterations and comparing them to the quantification of drivers in the future scenarios.

**Figure 1:**
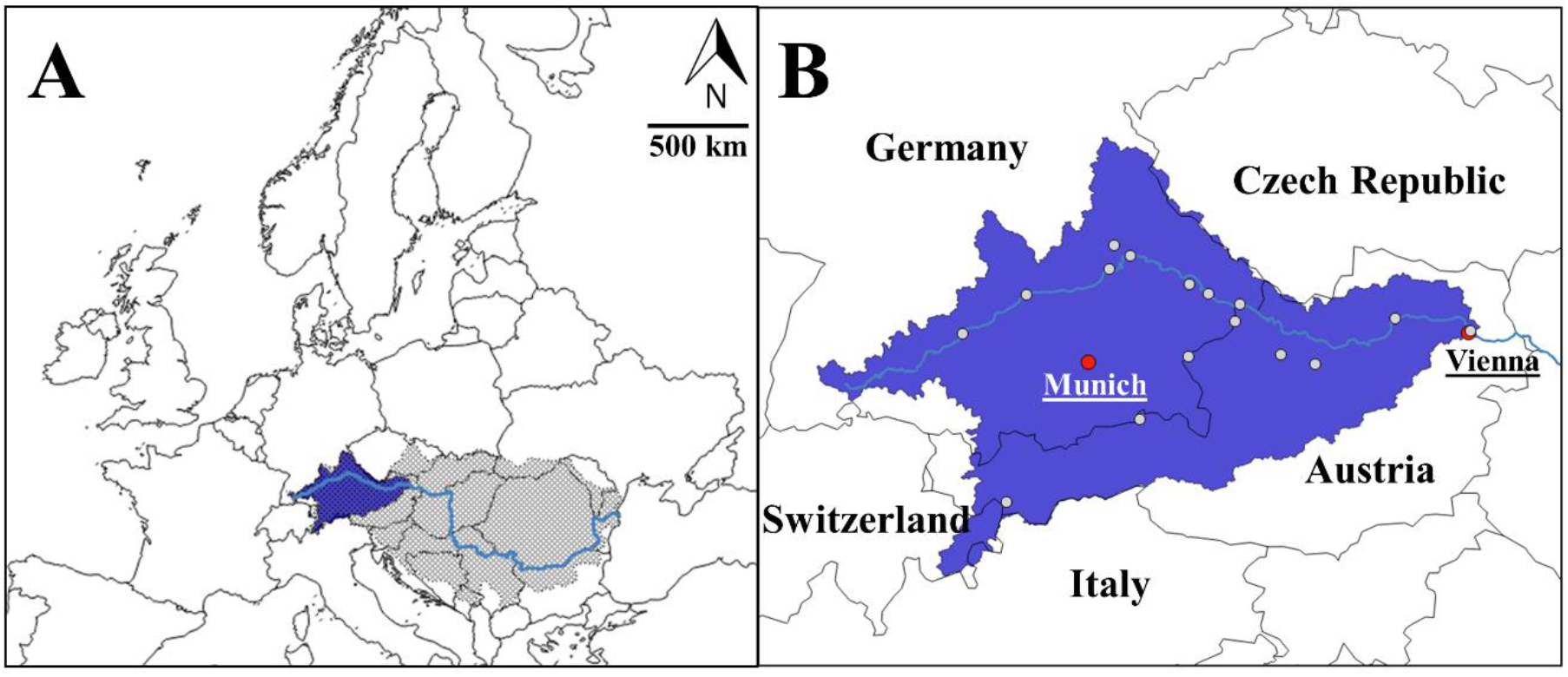
Overview of the study area. A) The location of the upper Danube River Basin within Europe. The light grey area indicates the whole Danube River Basin. In light blue the Danube main stem is shown. Dark blue indicates the study area. B) Study area in detail. Large red dots indicate the location of Munich and Vienna. Small grey dots show the distribution of the gaging stations we used to extrapolate discharge values across the study area (see “Hydrological data” for details).

We first used habitat suitability models (HSMs) to assess the current distribution of suitable habitats for 48 native fish species. We then used the current predicted fish habitat distributions (and species habitat relationships) as a baseline for climate niche factor analyses (CNFA; Rinnan & Lawler, 2019) to assess species vulnerability for historical alterations and future change scenarios. We hypothesised that (i) historical vulnerability estimates will be mainly driven by discharge-related environmental factors. We further expected that under future environmental change scenarios, the importance of discharge-related factors will play a lesser role than temperature-related environmental factors. In addition, we hypothesised (ii) that historical discharge alterations caused by the damming and channelisation affected riverine fish species stronger than the combined changes in climate and climate-driven flow characteristics would do in the future. We expect that this difference is indicated by overall higher historical vulnerability estimates than for future scenarios.

## Methods

### Study region

The study region is the upper Danube River Basin from its source in Germany’s Southwest up to the gaging station close to Vienna, Austria, covering 102.113 km^2^ and roughly 1000 km of the Danube River main stem. The upper Danube River Basin mainly covers parts of Germany and Austria (> 90%) and minor parts of Switzerland, Italy, and the Czech Republic (Fig. 1B). For subsequent analyses, we divided the study area into >18.000 sub-basins (Fig. 2A) and considered each sub-basin with at least one occurrence of a particular species as a sub-basin with said species presence (Fig. 2B).

**Figure 2:**
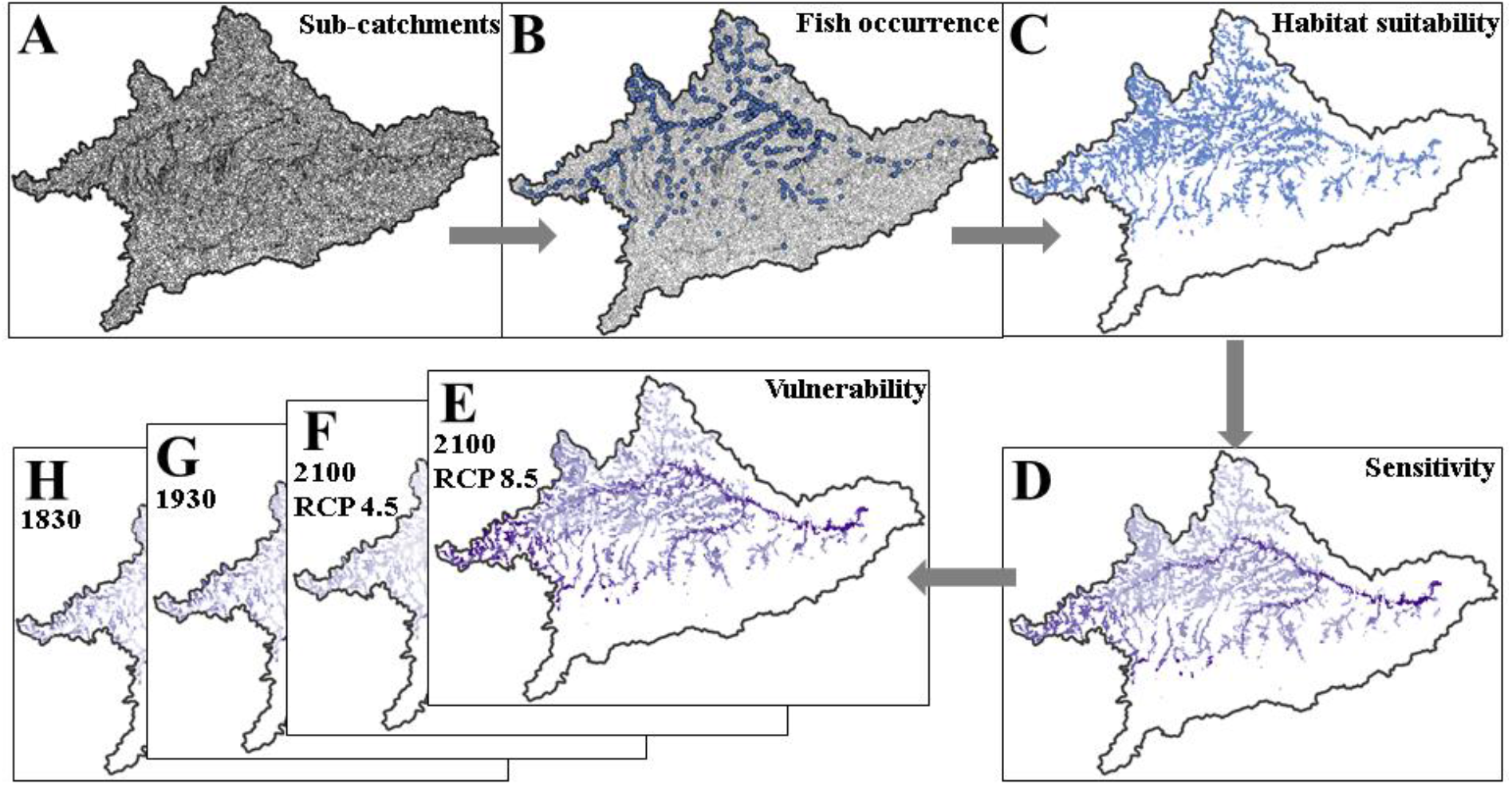
General workflow. A) The study area divided into 18.708 sub-basins with a mean area of 8 ± 8 km^2^. B) Spatial overlay of the fish occurrence data (here the common bream, *Abramis brama*) with the sub-basins in the study area. C) For each species occurring in at least ten sub-basins, present-day habitat suitability was calculated based on the sub-basins as modelling units and six environmental predictors (i.e. mean annual temperature, temperature annual range, mean monthly discharge, coefficient of variance of monthly discharge, mean northness, and roughness range), to obtain a range-wide estimate of the habitat suitability for a target species. D) A sub-basin-specific sensitivity estimate for each species within its potential distribution was calculated based on the average sensitivity of a given species (i.e. the average environmental conditions used by that species) compared to the average environmental conditions within a given sub-basin. For instance, a low sensitivity would be assigned to a sub-basin, where the environmental conditions match the average environmental conditions used by that species across the entire study area, and vice versa. E-H) For each species, a sub-basin specific vulnerability estimate for the different time intervals and climate change scenarios was calculated (for details see Departure and vulnerability analysis).

### Fish distribution data

We compiled fish species occurrence data from four different sources. For the German part of the upper Danube River Basin, we used the occurrence data collected by the Federal Ministries of Bavaria and Baden-Württemberg to comply with the EU Water Framework Directive. For the Austrian part of the upper Danube River Basin, we used occurrence data collected for the project “Improvement and Spatial extension of the European Fish Index” (EFI+, Pont et al., 2009). In addition to this large-scale monitoring data, we used occurrence data from Brunken et al. (2008). From Brunken et al. (2008) we only used data collected by acknowledged sources, such as universities and federal ministries for species listed in the data sources from Bavaria, Baden Württemberg or EFI+. We filtered all fish occurrence data for sampling dates between 1970 and 2016. Here, we included data from 2000 to 2016 as this resulted in an addition of almost 60% of high-quality fish records compared to 1970–2000. In total we gathered data for 48 native fish species (Tab. S1).

### Environmental data time-series

Hydrological and climate data were obtained from the German Federal Institute of Hydrology (BfG; Stanzel & Kling, 2018).

#### Hydrological data

Monthly hydrological data for 16 gaging stations within the study area (Fig. 1B) were obtained from the precipitation runoff model COSERO for a time interval from 1800 to 2100 (Stanzel & Kling, 2018). Historical discharge simulations from COSERO are based on gridded monthly temperature data and precipitation data available in the HISTALP data base (Auer et al., 2007; Chimani et al., 2013). Future discharge simulations are based on temperature and precipitation projections of Regional Climate Models (RCMs) from the EUR-11 ensemble of the EURO-CORDEX initiative (Jacob et al., 2014). We used future discharge simulations based on ten different global-regional climate model (GCM/RCM) combinations (i.e. CERFACS-CNRM-CM5/CCLM4-8-17, EC-EARTH/CCLM4-8-17, HadGEM2-ES/CCLM4-8-17, M-MPI-ESM-LR/CCLM4-8-17, EC-EARTH/RACMO22E, HadGEM2-ES/RACMO22E, EC-EARTH/HIRHAM5, IPSL-CM5A-MR/WRF331F, CERFACS-CNRM-CM5/ALADIN53, M-MPI-ESM-LR r2i1p1/REMO2009) and for the two Representative Concentration Pathways (RCP; van Vuuren et al., 2011) 4.5 and 8.5 (Stanzel & Kling, 2018). For further details on COSERO and its performance see Klein et al. (2011); Kling, Fuchs, et al. (2012); Kling, Lagler, et al. (2012). To create discharge estimates for each sub-basin, we extrapolated the modelled discharge values from the 16 gaging stations using a linear model of flow accumulation (number of grid cells contributing to a given stream grid cell) and monthly discharge (see also Kuemmerlen et al., 2014). From the entire time series of simulated monthly discharge data, we extracted two historic time intervals (1800-1830 and 1900-1930), a current time interval (1970-2000) and a future time interval (2070-2100) with two scenarios (RCP 4.5 and RCP 8.5). For each time interval and scenario, we calculated the coefficient of variance of monthly discharge and the mean annual discharge and for each sub-basin (see Tab. 1 for an overview of the raw values), which were later used as predictors for the habitat suitability modelling (HSM; only the current time interval) and CNFA.

**Table 1:** Summary statistics of predictors used for the HSM and CNFA analyses (static topography related predictors are not shown). For each time interval and scenarios, the mean value ± standard deviation, the median value with the first and third quartile, and the highest and the lowest value are shown. CoV = coefficient of variance, sd = standard deviation.

| Time interval | Predictor | Mean (± sd) | Median (1<sup>st</sup> and 3<sup>rd</sup> quartile) | Highest value | Lowest value |
| --- | --- | --- | --- | --- | --- |
| 1800 - 1830 | CoV of monthly discharge | 2.23 (0.93) | 2.70 (1.62; 2.98) | 3.06 | 0.25 |
|  | Mean annual discharge (m<sup>3</sup>/s) | 322.31 (747.03) | 67.7 (61.60; 120.93) | 3251.50 | 8.28 |
|  | Temperature annual range (°C) | 25.99 (2.64) | 26.57 (26.11; 27.33) | 28.37 | 3.31 |
|  | Mean annual temperature (°C) | 6.69 (2.04) | 7.13 (6.72; 7.79) | 8.52 | -3.01 |
| 1900 - 1930 | CoV of monthly discharge | 2.28 (0.96) | 2.75 (1.61; 3.06) | 3.14 | 0.27 |
|  | Mean annual discharge (m<sup>3</sup>/s) | 313.87 (744.37) | 61.50 (55.60; 113.42) | 3268.10 | 7.51 |
|  | Temperature annual range (°C) | 24.22 (2.63) | 24.80 (24.35; 25.65) | 26.51 | 3.10 |
|  | Mean annual temperature (°C) | 6.36 (2.02) | 6.83 (6.33; 7.49) | 8.07 | -3.28 |
| 1970 - 2000 | CoV of monthly discharge | 1.45 (0.55) | 1.74 (1.13; 1.87) | 1.91 | 0.21 |
|  | Mean annual discharge (m<sup>3</sup>/s) | 337.35 (741.17) | 85.40 (79.50; 137.73) | 3276.10 | 10.50 |
|  | Temperature annual range (°C) | 24.73 (2.63) | 25.30 (24.89; 26.13) | 27.10 | 3.16 |
|  | Mean annual temperature (°C) | 7.13 (2.03) | 7.67 (7.04; 8.29) | 8.78 | -2.56 |
| 2070 - 2100 RCP 4.5 | CoV of monthly discharge | 1.29 (0.50) | 1.55 (0.99; 1.68) | 1.72 | 0.20 |
|  | Mean annual discharge (m<sup>3</sup>/s) | 332.30 (741.58) | 80.40 (74.50; 132.36) | 3266.20 | 9.88 |
|  | Temperature annual range (°C) | 25.20 (2.58) | 25.77 (25.27; 26.62) | 27.52 | 3.21 |
|  | Mean annual temperature (°C) | 9.03 (2.04) | 9.51 (8.97; 10.20) | 10.82 | -0.44 |
| 2070 - 2100 RCP 8.5 | CoV of monthly discharge | 1.51 (0.67) | 1.84 (0.98; 2.06) | 2.13 | 0.22 |
|  | Mean annual discharge (m<sup>3</sup>/s) | 314.92 (741.35) | 64.30 (58.70; 113.26) | 3273.40 | 7.87 |
|  | Temperature annual range (°C) | 25.20 (2.53) | 25.87 (25.22; 26.56) | 27.62 | 3.23 |
|  | Mean annual temperature (°C) | 10.58 (1.92) | 11.02 (10.56; 11.65) | 12.35 | 0.95 |

#### Temperature data

To be consistent with the discharge model COSERO, we used the same monthly temperature data (i.e. same historic data and the same regional climate models). COSERO is driven by temperature data, which was downscaled to 61 hydrological response units within the study area based on elevation (for details see Kling, Lagler, et al., 2012). We used this downscaled monthly climate data to calculate mean annual temperatures and temperature annual ranges for each sub-basin and each of the aforementioned four time intervals (1800-1830, 1900-1930, 1970-2000, and 2070-2100 with RCP 4.5 and RCP 8.5; see Tab. 1).

#### Topographical data

Topographical data was used for the present-day HSM, because topography-related predictors were found to be important when modelling habitat suitability of fish species in the upper Danube River Basin (Friedrichs-Manthey et al., 2020). Topography is not expected to change over the analysed 300 years’ time period, therefore we used topography related predictors only to obtain the best possible current predictions in our HSM approach and excluded them from the CNFA analyses. Topographical data was obtained from the EarthEnv topography dataset (Amatulli et al., 2018), providing topographical data on a 1 km^2^ resolution globally. From these, we calculated the mean northness and roughness range for each sub-basin.

### Habitat suitability modelling

To further increase spatial completeness of the compiled distribution data, we used habitat suitability models (Elith & Leathwick, 2009). HSMs use a statistical relationship between species occurrence data and environmental predictors to create range-wide predictions of habitat suitability for target species. We built models using the biomod2 package in R (R-Core-Team, 2013; Thuiller et al., 2009). We used weighted ensemble models (Marmion et al., 2009) consisting of five machine learning and regression algorithms (Artificial Neural Networks, Maximum Entropy, Generalized Linear Model, Generalized Additive Model and Multivariate Adaptive Regression Splines) that are widely applied within the HSM literature (Araujo & New, 2007; Merow et al., 2014). We only used species with at least ten unique occurrences (i.e., present in sub-basins) to model the potential habitat suitability (van Proosdij et al., 2016) and a fixed number of one third of all sub-basins as pseudo absences. As predictors we used six uncorrelated environmental variables from three different categories, which have shown to be appropriate to model habitat suitability of fish species in the upper Danube River Basin (Friedrichs-Manthey et al., 2020): hydrology - average annual discharge and coefficient of variance of monthly discharge, climate - mean annual temperature and temperature annual range, and topography - average northness and roughness range. We assigned proportional weights to all single models (e.g. single algorithms) before combining them to a final ensemble model for each species. The assignment of weights allows focusing on the best algorithm without discarding results from other algorithms completely (Araujo & New, 2007). Weights were assigned according to the True Skill Statistic (TSS, Allouche et al., 2006). TSS values range between −1 and +1, with values around zero indicating that a model is not better than random and values of +1 indicating a perfect fit. The final ensemble models were evaluated using TSS by means of data splitting: we used ten separate model runs, where 70% of the data was used for calibration and 30% for model validation. Predicted habitat suitability was transformed to a binary presence/absence information (Fig. 1D; from here on called suitable habitats of a species) using a species-specific cut-off value, which minimises the absolute difference between sensitivity (i.e., true positive rate: how well a model depicts the true known presences of a certain species) and specificity (i.e., true negative rate: how well a model depicts the randomly created pseudo-absences). Note that the model sensitivity in the HSM validation is disconnected from a sensitivity estimate used in the CNFA analyses, described below.

### Departure and vulnerability analysis

We used the CNFA package in R (Rinnan & Lawler, 2019) to calculate average environmental sensitivity estimates for all species, species-specific departures between the time intervals, and species specific vulnerability estimates for each time interval except for the current point in time. The average environmental sensitivity reflects the average degree of specialisation of a species for each environmental predictor (Rinnan & Lawler, 2019). These environmental sensitivity values are always positive and interpreted in comparison among analysed species. The higher the sensitivity of a species, the smaller the estimated environmental niche of that species given the predictors used in relation to other species. In a second step, we calculated the species-specific departure (i.e. the environmental distance) between the current time interval and the two historical time intervals and two future scenarios for each sub-basin. Departure is defined as a measure of change between baseline habitat conditions (here the present day conditions) and historic or future habitat conditions (Rinnan & Lawler, 2019). The departure estimate is always positive and has no upper limit. In total, we calculated 22 departure estimates for each species (2 historic, and 2 future * 10 RCMs). As the ten RCMs were highly variable and consequently produced a large variety in departure estimates, we used the median to create one future departure layer per species and RCP, combining all ten RCM models. All calculations are relative to the present, modelled habitat suitability of a species. Finally, vulnerability estimates are a combination of environmental sensitivity and departure. The species-specific sensitivity estimate weights the overall departure estimate of a given location. Consequently, high sensitivity values and high departure estimates result in a predicted high vulnerability. A standalone vulnerability estimate has no meaning but does so when compared to species or within species for different time intervals. We calculated the vulnerability for each sub-basin given the species that have a present-day predicted suitable habitat for the two historical time intervals and the two future scenarios (Fig. 1 E-H). Again, we calculated the ten future climate models separately and then combined them to one vulnerability layer for each species and future scenario using the median. We compared the four resulting mean vulnerability layers using the ‘modOverlap’ function in the R package fuzzySim (Barbosa, 2015) and calculated Schoeners’D (Warren et al., 2008), which can range from 0 (no overlap) to 1 (total overlap), between each time interval and the two future scenarios.

## Results

### SDM performance

The TSS values for all species ranged between 0.40 for species with >400 initial occurrences (*Salmo trutta*) to 0.99 for species with <20 occurrences (i.e. *Telestes souffia*), indicating a good to very good model performance for all species (Bean et al., 2012; for details see Tab. S1).

### Departure

Relative to the suitable habitats modelled for the current point in time (1970-2000), the median departure values, i.e. the environmental distance, for the environmental predictors showed large differences between the analysed time intervals and scenarios. The environmental departure of the coefficient of annual discharge was similar between the two historic time intervals (1800-1830: 0.58, 0.49 and 0.69; 1900-1930: 0.60, 0.51 and 0.72; median, 1^st^ and 3^rd^ quartile, respectively) and more than two times higher than the median departure for the RCP 4.5 scenario (0.23, 0.19 and 0.27). Compared to the RCP 4.5 scenario, the departure increased for the RCP 8.5 scenario (0.39, 0.32 and 0.47), but remained lower than the departure values for the historic time intervals (Fig. 3, light blue box-plots).

**Figure 3:**
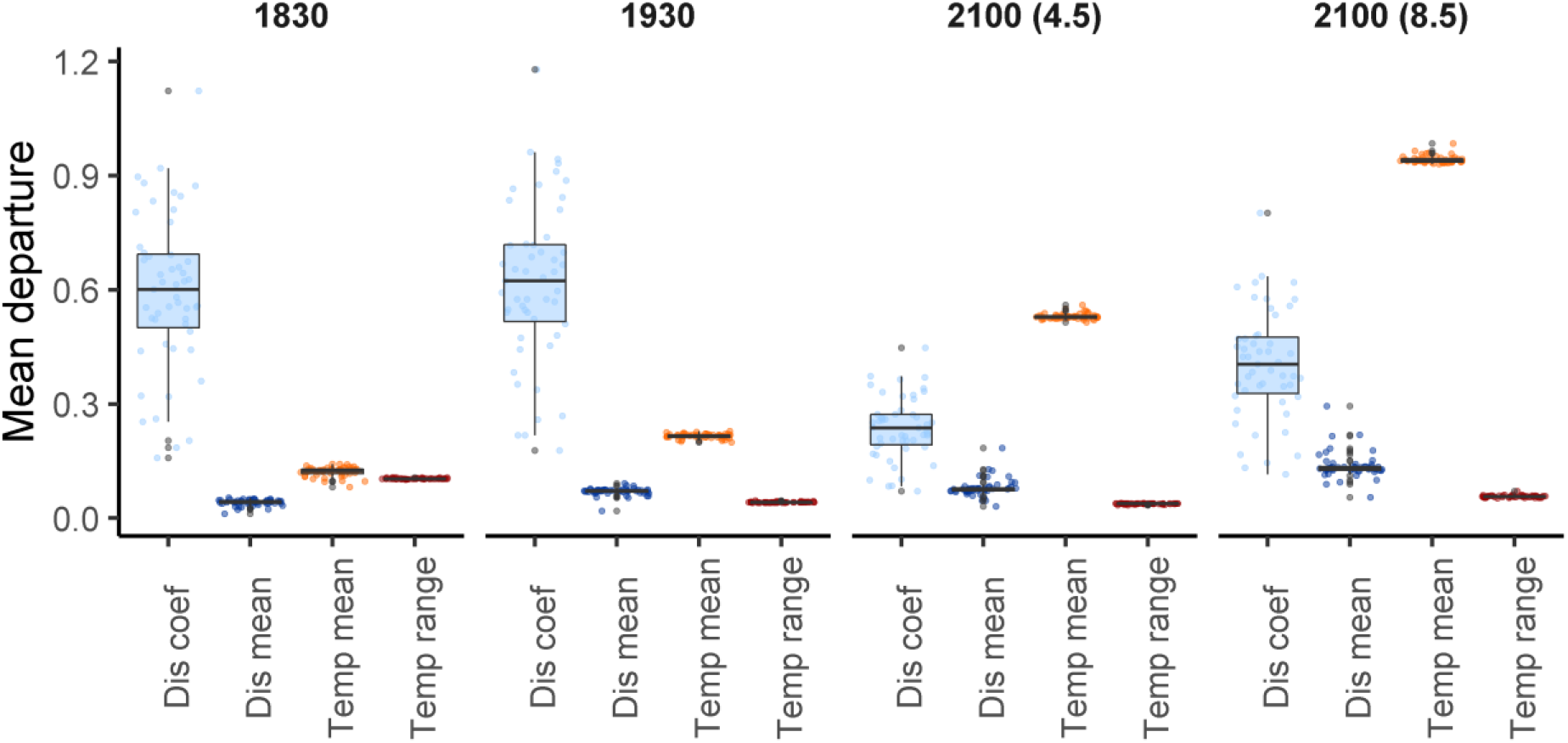
Mean departure values for the four environmental predictors used in the CNFA for 48 native fish species in the upper Danube River Basin. Departure is always measured as the distance between the current point in time (1970-2000) and the respective time interval or scenario. Dis coef = Coefficient of variance of annual discharge, Dis mean = Mean annual discharge, Temp mean = Mean annual temperature, Temp range = Range of mean monthly temperatures.

The median departure for the annual mean discharge constantly increased from the historic time interval 1800-1830 to the historic time interval 1900-1930 and to the future scenarios, with its highest values for the future RCP 8.5 scenario (0.13, 0.13 and 0.14: Fig. 3, dark blue box-plots). We found the same pattern for the median departure for the annual mean temperature: the highest departure was calculated for the future RCP 8.5 scenario with a median value of 0.94 (0.94 and 0.94, Fig. 3, orange box-plots). We found the opposite pattern for the median departure for the annual temperature range. The median departure was highest in the historic time interval 1800-1830 (0.1, 0.1 and 0.1) and lowest for the future RCP 4.5 scenario (0.04, 0.04 and 0.04) and only slightly increased for the future RCP 8.5 scenario (0.06, 0.06 and 0.06, Fig. 3, red box-plots).

### Vulnerability

Median vulnerability for all 48 native fish species was lowest for the RCP 4.5 scenario (0.55, 0.51 and 0.66; median, 1^st^ and 3^rd^ quartile, respectively) and almost doubled for the RCP 8.5 scenario (1.02, 0.91 and 1.16; Fig. 4). The two historic time intervals ranged in between the future scenarios (distant historic: 0.60, 0.47 and 0.74; near historic: 0.65, 0.52 and 0.79; Fig. 4).

**Figure 4:**
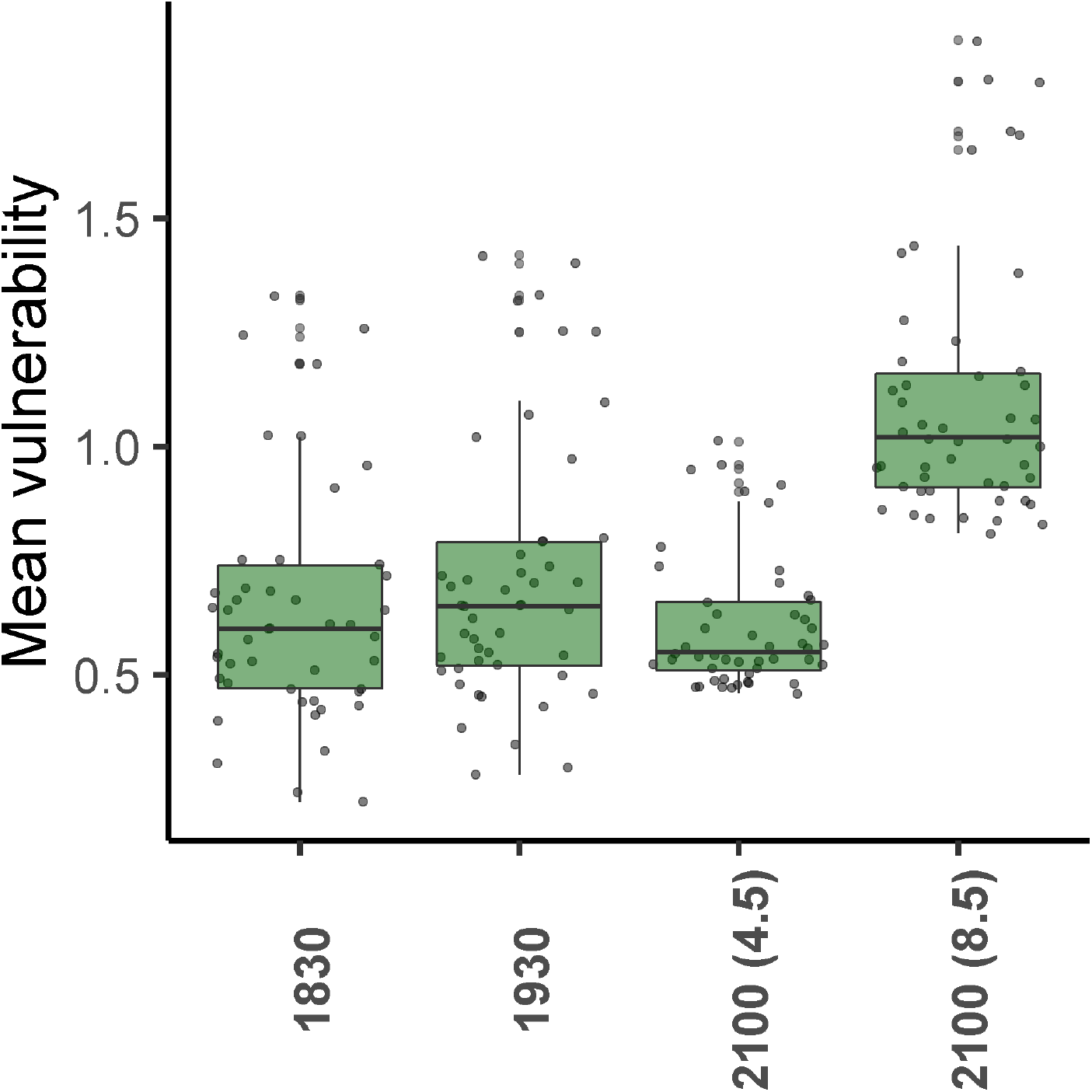
Mean vulnerability estimates for 48 native fish species in the upper Danube River Basin. Note: The y-axis shows species mean vulnerability which is calculated as the mean value across all sub-basins in the predicted distribution range for each of the 48 species.

The comparison of similarity (Schoeners’D; Warren et al., 2008) between predicted mean vulnerability estimates showed that similarity between scenarios was highest for the historical time intervals (0.99; Fig. 5 B). The two future scenarios showed a high similarity as well (0.96). Similarity was lowest when comparing any historical time interval with any future scenario (0.88 for both historical scenarios against future RCP 4.5; Fig. 5 B).

**Figure 5:**
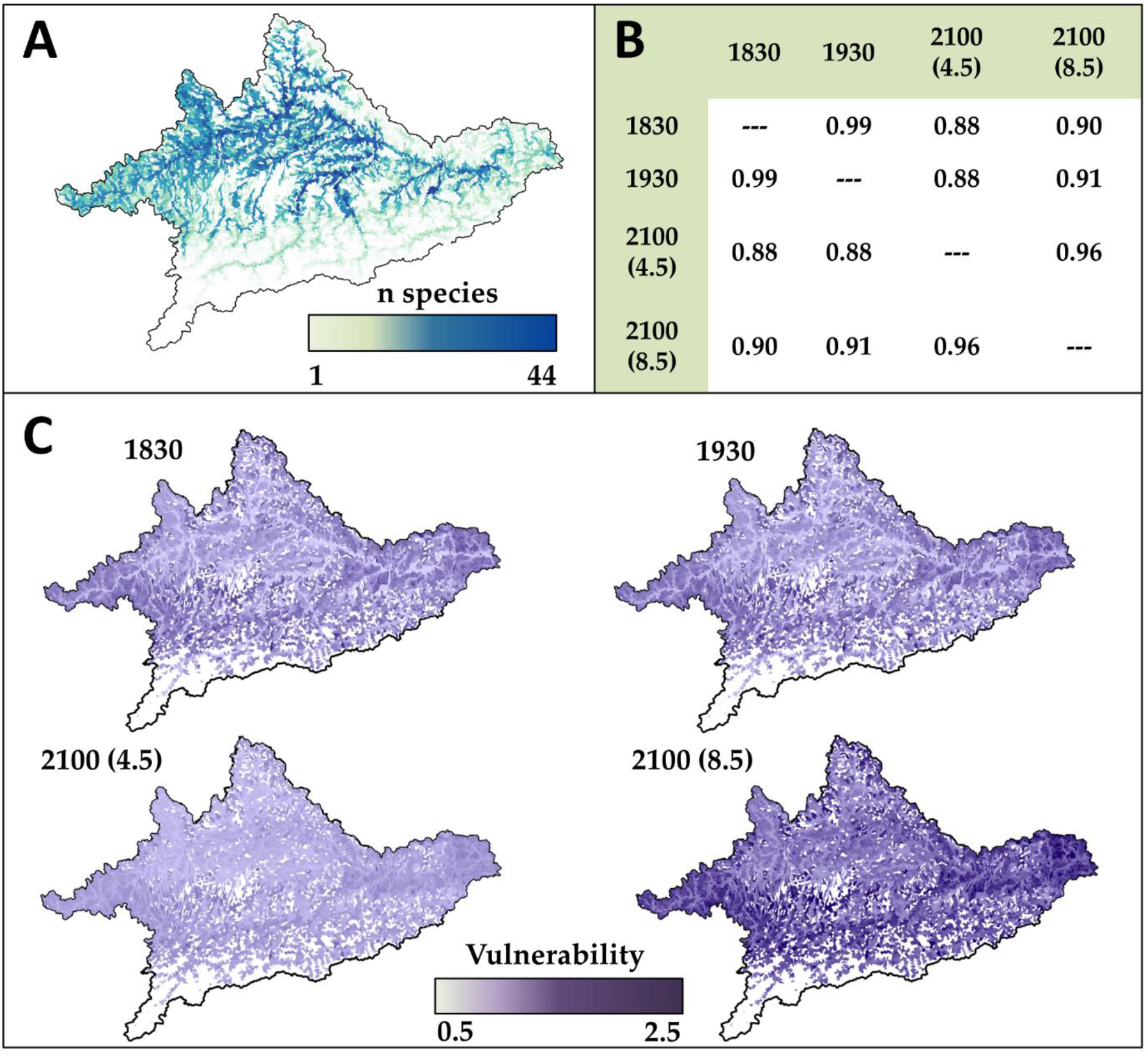
A) Number of species predicted to find suitable habitats within a sub-basin. B) Similarity (Schoeners’D; Warren et al., 2008) between predicted vulnerability estimates for the two historic time intervals and the two future scenarios. C) Mean vulnerability for each sub-basin for the two historic time intervals and the two future scenarios, all relative to the present distribution range.

## Discussion

In our study we showed that the drivers of species-specific vulnerability shifted in their importance from hydrological predictors in the past to climatic predictors in the future. Resulting vulnerability estimates were in terms of magnitude similar between the historical time intervals and the moderate RCP 4.5 scenario. However, for the more severe RCP 8.5 scenario we identified an almost two-fold increase in vulnerability estimates compared to the RCP 4.5 scenario.

### Fish species vulnerability in a historical context

Humans started to regulate the upper Danube River Basin as early as at the end of the 16^th^ century (Jungwirth et al., 2014). Flood control measures and improvement of inland navigation, especially for the Austrian Danube River Basin, were the main force driving river alteration. By the end of the 19^th^ century, flood control and channelization resulted in a loss of 15% of river length for the German and Austrian Danube main stem (Jungwirth et al., 2014). Main stem embankment reached its maximum at the beginning of the 20th century. To date, more than 90% of all floodplains have been disconnected along the German part of the Danube (Brunotte et al., 2009), while the shoreline of the upper Danube mainstem is to more than 90% embanked (Jungwirth et al., 2014). Those anthropogenic changes have influenced the seasonal patterns of the discharge regime in the upper Danube River Basin with an increase of mean monthly flows in winter months and a decrease in the summer months, whereas the overall mean annual flow remained constant over the last 100 years (Klein et al., 2011). We found that these historically documented habitat alterations did not result in pronounced differences of vulnerability estimates for the analysed fish species for the two historical time intervals (1800-1830 and 1900-1930; Fig. 4, Fig. 5 C). This finding seems contradictory, since the more pristine upper Danube River Basin around 1800-1830 was environmentally more distant to the current point in time than the more regulated Danube River Basin around 1900-1930. However, flood regulation and river channelisation measures during the 19^th^ century were mainly made of wood and, therefore, were not durable against annual ice floods and had to be renewed almost every year (Hohensinner et al., 2013). Therefore, their effect on river discharge regimes was likely low as supported by our results. Those measures did neither affect annual variability in discharge nor monthly discharge, or not in a way that would have been captured by our hydrological predictors with a temporal resolution of a month. Discharge data on a daily resolution could potentially capture a higher variability than our monthly predictors, however, those predictors are currently not available and we can only conclude that the flood regulation and river channelisation measures applied in the beginning of the 19^th^ century had a relatively low effect on our vulnerability estimates.

In contrast, we found a pronounced change in hydrological conditions from the current point in time to 1900-1930 mainly caused by a change in the variability of discharge (Fig. 3). Interestingly, after channelisation was mostly completed at the beginning of the 20th century (at least for the main stem), damming started in many tributaries and in the Danube main stem in the upper Danube River Basin (Bettzieche, 2010; Habersack et al., 2016; Jungwirth et al., 2014). To date, countless, mainly small (< 1MW) hydropower plants exist in the upper Danube River Basin (Habersack et al., 2016). In the Danube main stem in Germany and Austria more than 70% of the river length is dammed (Jungwirth et al., 2014) and free-flowing river stretches disappeared (Brinker et al., 2018; Duarte et al., 2020; Schiemer & Spindler, 1989). Our predictors and, consequently, our models picked up on those range-wide hydrological changes, resulting in a dramatic decrease in variability of monthly discharge from 1900-1930 to the current point in time. However, model results showed similar average vulnerability estimates for fish species for both historical time intervals and the climate change scenario under RCP 4.5 (Fig. 4). This finding leads to the assumption that historical discharge alterations induced by dams have resulted in comparable pressures for native species as future changes under RCP 4.5 will do. Since we found a change in the main driver of vulnerability estimates between 1900-1930 and 2070-2100, i.e. from variability in discharge to mean annual temperature, respectively, we can assume that, ecologically, future climate change will impose a different threat to native fish species than the historical discharge-driven alterations, at least for the moderate RCP 4.5 scenario. Regarding the more extreme RCP 8.5 scenario, which tracks current CO2-emissions best (Schwalm et al., 2020), we observed a pronounced increase in overall vulnerability estimates compared to the two historical time intervals and the RCP 4.5 scenario (Fig. 4). This increase was mainly driven by an overall +4 °C increase in mean annual temperature (compared to the current status and a +2 °C mean increase compared to the RCP 4.5; Tab. 1). Interestingly, the variability of monthly discharge returned as an essential driver of vulnerability estimates for RCP 8.5. Hence, our results indicate that future climate change would cause temperature-driven flow alterations comparable to historical anthropogenic alterations. Considering that COSERO does not account for new dams planned to be built, the cause for these discharge alternations are a result of changes in climate and precipitation only. Consequences for the native fish species in the upper Danube River Basin under RCP 8.5 would be a significant temperature increase and additional hydrological pressures, similar in magnitude to what they have already experienced historically.

### Practical implications

For some organism groups, such as benthic invertebrates (Durance & Ormerod, 2009) or marine fish (Roberts et al., 2017), a reduction of environmental pressures can promote the resilience towards anticipated climatic pressures. For instance, Durance and Ormerod (2009) showed that for benthic invertebrate communities in small streams, expected changes in species community due to warming waters over an 18 year period were buffered by a steadily increasing water quality over the same time period. Hence in simple terms, a decrease in one pressure balanced an increase in another pressure. Our results indicate an analogy for the upper Danube River Basin and its fish community. The overall environmental distance caused by the reduction of monthly discharge variability is similar to the overall environmental distance that would be expected under future climate change scenarios, which is induced by increased mean annual temperatures.

For the upper Danube River Basin only a few species, mainly anadromous sturgeons, went regionally extinct when comparing the current fish community with the community around 1800 (Friedrich, 2018; Hensel & Holcík, 1997). For the sturgeons, this regional extinction is mainly caused by large dams which act as migration barriers and poaching in the lower Danube regions (Jungwirth et al., 2014). For other fish species “only” a pronounced change in relative abundance after damming was observed (Galik et al., 2015; Schmutz et al., 2013). This observation indicates that the historical fish community itself is still present in the upper Danube River Basin, an important precondition for effective fish species conservation in river ecosystems (Stoll et al., 2014). To relieve the environmental pressure induced through hydrological alterations, conservation measures such as floodplain rehabilitation have shown to be very effective for fish communities (Ramler & Keckeis, 2019; Roni et al., 2008). In the upper Danube River Basin, roughly 25% of the historically available and nowadays unconnected floodplain area has a good potential for rehabilitation measures (Hein et al., 2016). Additionally, in view of the increasing temperature pressure, the upper Danube River Basin with its many headwater regions might offer a suitable temperature refuge for sensitive fish species (Isaak et al., 2016).

While our study provides a generally promising outlook, we acknowledge that it is also a result of the “survivorship bias” (Budd & Mann, 2018). Using monitoring data from 1970 to 2016, we excluded species that went already regionally extinct before 1970 from our analyses. Additionally, using only species with more than ten occurrence points, we excluded range-restricted species that are either hard to detect or less abundant (Cruickshank et al., 2016). Furthermore, the species-environment relationship that we have used to model habitat suitability might be incomplete, since species might not reach all suitable habitats due to, i.e. migration barriers. Extinction and low abundance can both be a result of historic discharge alterations. By excluding those species, and the possibility that some species-environment relationships are incomplete, we likely underestimated the effect of historic discharge alterations in the upper Danube River Basin.

### A conservative approach to reduce uncertainty

In this study, we modelled how native fish species in the upper Danube River Basin were affected by historic environmental alterations and how they would be affected by future climate change within their current distribution ranges. We used HSMs to fill monitoring gaps of the current distribution of species, but we did not use them to assess potential changes in their spatial distribution neither historically nor in the future, as it is usually done (Ehrlen & Morris, 2015; McMahan et al., 2020; Radinger et al., 2017). In contrast, we analysed the environmental conditions that drive the habitat suitability of the species at different points in time. We decided to use this more conservative approach, because it is well understood that any prediction based on HSMs comes at the cost of uncertainty, especially when a model is transferred to new environments or time frames (Werkowska et al., 2017; Yates et al., 2018). In this sense, our approach has the advantage that we can be more certain about the expected changes in the future within the current distribution range of a species, at least regarding the predictors we used. Therefore, our results have the potential to provide guidance towards future conservation actions and conservation management (Schuwirth et al., 2019).

## Acknowledgements

This work was funded by the German Federal Ministry of Education and Research (BMBF) within the “GLANCE” project (Global Change Effects in River Ecosystems; 01 LN1320A). We further acknowledge funding by the European Union’s Horizon 2020 Research and Innovation Programme grant number 642317. SD acknowledges funding by the Leibniz Competition (J45/2018). SDL has received funding from the European Union’s Horizon 2020 Research and Innovation Programme under the Marie Skłodowska-Curie Grant agreement No 748625. FB and TH acknowledge support from the Christian Doppler Society – CD Laboratory for meta ecosystem dynamics in riverine landscapes. MFM wants to thank the IGB for supporting early career scientists with care obligations during the Corona pandemic. We thank the EU projects Biofresh (Contract No 226874) and EFI+ (Contract No 044096) for support with occurrence data. Support by Martin Winkler in data handling from the Fischartenatlas is greatly acknowledged.

## Notes

### Competing Interest Statement

The authors have declared no competing interest.

