## Supplementary Table S1 for "300 years of change for native fish species in the upper Danube River Basin – historical flow alterations versus future climate change"

Supplementary Table S1: Details on the number of initial occurrences, the number of sub-basins considered as suitable for a species after the HSM, the TSS value for the ensemble model, the mean (± standard deviation) sensitivity value, and mean (± standard deviation) vulnerability estimates for the three time intervals and two future scenarios for all 48 modelled native fish species.

| Species | Occurrences | n sub-basins after HSM | TSS (ensemble) | Mean sensitivity | 1830 mean vulnerability | 1930 mean vulnerability | RCP 4.5 mean vulnerability | RCP 8.5 mean vulnerability |
| --- | --- | --- | --- | --- | --- | --- | --- | --- |
| *Abramis brama* | 526 | 5302 | 0.54 | 0.67 ± 0.34 | 0.48 ± 0.19 | 0.51 ± 0.19 | 0.47 ± 0.11 | 0.85 ± 0.20 |
| *Alburnoides bipunctatus* | 480 | 5411 | 0.53 | 0.65 ± 0.31 | 0.47 ± 0.17 | 0.50 ± 0.17 | 0.48 ± 0.11 | 0.84 ± 0.20 |
| *Alburnus alburnus* | 637 | 6419 | 0.54 | 0.68 ± 0.37 | 0.53 ± 0.20 | 0.56 ± 0.20 | 0.48 ± 0.13 | 0.87 ± 0.23 |
| *Alburnus mento* | 11 | 248 | 0.99 | 2.27 ± 1.12 | 1.26 ± 0.34 | 1.32 ± 0.37 | 0.96 ± 0.02 | 1.80 ± 0.41 |
| *Aspius aspius* | 154 | 3697 | 0.64 | 0.95 ± 0.48 | 0.33 ± 0.13 | 0.38 ± 0.13 | 0.53 ± 0.16 | 0.95 ± 0.24 |
| *Ballerus sapa* | 45 | 2615 | 0.88 | 0.90 ± 0.60 | 0.22 ± 0.11 | 0.28 ± 0.12 | 0.53 ± 0.16 | 0.93 ± 0.28 |
| *Barbatula barbatula* | 1037 | 6841 | 0.47 | 1.99 ± 1.20 | 1.18 ± 0.31 | 1.25 ± 0.33 | 0.90 ± 0.20 | 1.68 ± 0.38 |
| *Babus barbus* | 674 | 5286 | 0.54 | 0.65 ± 0.29 | 0.40 ± 0.15 | 0.43 ± 0.15 | 0.46 ± 0.10 | 0.81 ± 0.18 |
| *Blicca bjoerkna* | 251 | 5303 | 0.62 | 0.94 ± 0.59 | 0.60 ± 0.24 | 0.65 ± 0.24 | 0.56 ± 0.16 | 1.02 ± 0.28 |
| *Carassius carassius* | 129 | 5638 | 0.62 | 0.98 ± 0.53 | 0.75 ± 0.25 | 0.80 ± 0.26 | 0.60 ± 0.14 | 1.12 ± 0.27 |
| *Carassius gibelio* | 232 | 6338 | 0.56 | 0.85 ± 0.53 | 0.68 ± 0.25 | 0.72 ± 0.26 | 0.56 ± 0.16 | 1.03 ± 0.29 |
| *Chondrostoma nasus* | 479 | 5377 | 0.55 | 0.69 ± 0.32 | 0.42 ± 0.16 | 0.45 ± 0.16 | 0.47 ± 0.11 | 0.84 ± 0.20 |
| *Cobitis elongatoides* | 32 | 1924 | 0.87 | 1.84 ± 1.30 | 1.24 ± 0.38 | 1.33 ± 0.4 | 0.92 ± 0.26 | 1.69 ± 0.46 |
| *Cottus gobio* | 1401 | 6397 | 0.46 | 1.94 ± 0.94 | 1.18 ± 0.34 | 1.25 ± 0.35 | 0.88 ± 0.18 | 1.65 ± 0.35 |
| *Cyprinus carpio* | 550 | 5876 | 0.56 | 0.76 ± 0.41 | 0.61 ± 0.22 | 0.65 ± 0.22 | 0.53 ± 0.13 | 0.97 ± 0.24 |
| *Esox lucius* | 900 | 5540 | 0.52 | 0.68 ± 0.34 | 0.52 ± 0.19 | 0.55 ± 0.19 | 0.48 ± 0.11 | 0.88 ± 0.19 |
| *Eudontomyzon mariae* | 17 | 656 | 0.97 | 0.64 ± 0.25 | 0.43 ± 0.15 | 0.46 ± 0.14 | 0.48 ± 0.08 | 0.86 ± 0.15 |
| *Gasterosteus aculeatus* | 566 | 5379 | 0.57 | 0.78 ± 0.35 | 0.65 ± 0.24 | 0.70 ± 0.25 | 0.54 ± 0.11 | 1.01 ± 0.22 |
| *Gobio gobio* | 1037 | 5735 | 0.52 | 1.04 ± 0.63 | 0.75 ± 0.24 | 0.79 ± 0.25 | 0.62 ± 0.16 | 1.13 ± 0.30 |
| *Gymnocephalus baloni* | 33 | 2494 | 0.85 | 1.05 ± 0.75 | 0.58 ± 0.28 | 0.65 ± 0.28 | 0.63 ± 0.18 | 1.15 ± 0.31 |
| *Gymnocephalus cernua* | 120 | 6322 | 0.61 | 0.84 ± 0.54 | 0.66 ± 0.24 | 0.71 ± 0.25 | 0.57 ± 0.16 | 1.06 ± 0.29 |
| *Gymnocephalus schraetser* | 38 | 2488 | 0.88 | 1.29 ± 0.64 | 0.53 ± 0.28 | 0.59 ± 0.28 | 0.66 ± 0.15 | 1.16 ± 0.20 |
| *Hucho hucho* | 126 | 3715 | 0.65 | 0.87 ± 0.39 | 0.31 ± 0.13 | 0.35 ± 0.13 | 0.51 ± 0.10 | 0.92 ± 0.18 |
| *Lampetra planeri* | 68 | 3700 | 0.77 | 1.44 ± 0.84 | 1.02 ± 0.27 | 1.07 ± 0.28 | 0.78 ± 0.18 | 1.44 ± 0.34 |
| *Leucaspius delineatus* | 105 | 4355 | 0.74 | 1.37 ± 0.85 | 0.96 ± 0.28 | 1.02 ± 0.30 | 0.73 ± 0.17 | 1.38 ± 0.33 |
| *Leuciscus idus* | 252 | 6035 | 0.59 | 0.77 ± 0.47 | 0.54 ± 0.20 | 0.58 ± 0.20 | 0.52 ± 0.15 | 0.93 ± 0.26 |
| *Leuciscus leuciscus* | 866 | 5309 | 0.55 | 0.77 ± 0.42 | 0.58 ± 0.20 | 0.62 ± 0.20 | 0.53 ± 0.13 | 0.95 ± 0.23 |
| *Lota lota* | 409 | 6375 | 0.51 | 0.62 ± 0.36 | 0.49 ± 0.17 | 0.52 ± 0.18 | 0.47 ± 0.11 | 0.84 ± 0.20 |
| *Misgurnus fossilis* | 48 | 3108 | 0.80 | 1.16 ± 0.55 | 1.02 ± 0.26 | 1.10 ± 0.26 | 0.74 ± 0.12 | 1.42 ± 0.22 |
| *Perca fluviatilis* | 905 | 6727 | 0.48 | 0.73 ± 0.41 | 0.61 ± 0.19 | 0.65 ± 0.20 | 0.53 ± 0.12 | 0.96 ± 0.22 |
| *Phoxinus phoxinus* | 685 | 7585 | 0.47 | 2.56 ± 1.32 | 1.32 ± 0.39 | 1.40 ± 0.41 | 1.01 ± 0.21 | 1.89 ± 0.41 |
| *Proterorhinus marmoratus* | 40 | 1284 | 0.91 | 1.33 ± 0.81 | 0.46 ± 0.26 | 0.53 ± 0.27 | 0.63 ± 0.17 | 1.13 ± 0.30 |
| *Rhodeus amarus* | 314 | 5673 | 0.62 | 0.80 ± 0.49 | 0.60 ± 0.24 | 0.64 ± 0.24 | 0.53 ± 0.15 | 0.96 ± 0.27 |
| *Romanogobio vladykovi* | 99 | 5493 | 0.69 | 1.20 ± 0.67 | 0.74 ± 0.33 | 0.79 ± 0.33 | 0.66 ± 0.18 | 1.19 ± 0.32 |
| *Rutilus meidingeri* | 37 | 2575 | 0.79 | 0.72 ± 0.28 | 0.47 ± 0.21 | 0.51 ± 0.21 | 0.52 ± 0.10 | 0.91 ± 0.19 |
| *Rutilus rutilus* | 1131 | 6241 | 0.49 | 0.85 ± 0.51 | 0.68 ± 0.21 | 0.72 ± 0.22 | 0.56 ± 0.14 | 1.04 ± 0.26 |
| *Salmo trutta* | 2249 | 9168 | 0.40 | 2.23 ± 1.18 | 1.33 ± 0.39 | 1.42 ± 0.41 | 0.95 ± 0.20 | 1.80 ± 0.39 |
| *Salvelinus umbla* | 30 | 2765 | 0.85 | 1.02 ± 0.52 | 0.91 ± 0.26 | 0.97 ± 0.27 | 0.67 ± 0.16 | 1.28 ± 0.30 |
| *Sander lucioperca* | 242 | 6282 | 0.57 | 0.68 ± 0.37 | 0.55 ± 0.21 | 0.59 ± 0.22 | 0.49 ± 0.13 | 0.90 ± 0.23 |
| *Scardinius erythrophthalmus* | 437 | 5765 | 0.56 | 0.80 ± 0.43 | 0.66 ± 0.23 | 0.70 ± 0.24 | 0.55 ± 0.13 | 1.02 ± 0.25 |
| *Silurus glanis* | 179 | 6868 | 0.58 | 0.74 ± 0.37 | 0.64 ± 0.24 | 0.69 ± 0.24 | 0.54 ± 0.12 | 1.00 ± 0.24 |
| *Squalius cephalus* | 1463 | 5981 | 0.49 | 0.97 ± 0.57 | 0.72 ± 0.22 | 0.76 ± 0.23 | 0.60 ± 0.15 | 1.10 ± 0.27 |
| *Telestes souffia* | 18 | 347 | 0.99 | 0.82 ± 0.32 | 0.24 ± 0.13 | 0.30 ± 0.13 | 0.50 ± 0.10 | 0.90 ± 0.19 |
| *Thymallus thymallus* | 860 | 5718 | 0.58 | 0.78 ± 0.41 | 0.51 ± 0.17 | 0.54 ± 0.17 | 0.51 ± 0.11 | 0.91 ± 0.20 |
| *Tinca tinca* | 636 | 5042 | 0.52 | 0.87 ± 0.50 | 0.69 ± 0.22 | 0.74 ± 0.23 | 0.57 ± 0.13 | 1.05 ± 0.25 |
| *Vimba vimba* | 115 | 5067 | 0.64 | 1.01 ± 0.74 | 0.64 ± 0.31 | 0.69 ± 0.31 | 0.59 ± 0.18 | 1.06 ± 0.33 |
| *Zingel streber* | 56 | 2844 | 0.78 | 0.76 ± 0.28 | 0.41 ± 0.24 | 0.46 ± 0.24 | 0.49 ± 0.10 | 0.88 ± 0.19 |
| *Zingel zingel* | 30 | 1275 | 0.94 | 1.66 ± 0.63 | 0.44 ± 0.34 | 0.54 ± 0.35 | 0.70 ± 0.13 | 1.23 ± 0.25 |
